# Platelet-dependent clearance of uropathogenic *Escherichia coli* directly drives sepsis-induced thrombocytopenia in a mouse model

**DOI:** 10.1101/2025.05.23.655700

**Authors:** Nanna Johnsen, Mette G. Christensen, Laura V. Sparsø, Emil H. Lambertsen, Thomas Corydon, Helle Praetorius

## Abstract

Thrombocytopenia is a distinct negative prognostic marker in sepsis, a trait associated with the procoagulatory state of severe infection. However, thrombocytes have transcended to encompass a modulatory role in the immune response and as pathogen scavengers. In a murine model of urosepsis, we observed a substantial drop (40%) in circulating thrombocytes already 30 minutes after the introduction of uropathogenic *Escherichia coli* (UPEC) and a concomitant transient increase in both platelet factor 4 release and thrombin-antithrombin complexes. This reduction in thrombocytes was timely associated with a reduction in circulating UPEC. By imaging flow cytometry, we visualized that eGFP-expressing UPEC was instantly bound to circulating thrombocytes, leading to the immediate removal of thrombocyte-UPEC complexes and a 95% reduction in bacterial load within 10 minutes. We demonstrate that thrombocyte-UPEC complexes are cleared primarily through the liver, engaging the sinusoidal endothelial cells. The majority of the UPEC are recovered in the liver, with minimal contribution from intravascular bacterial damage or lysis. The thrombocyte-dependent clearance system has a maximal capacity, and overload markedly challenges the intravascular UPEC-clearance. The data strongly suggest that circulating thrombocytes constitute the most important cell type for fast scavenging and clearance of invading bacteria during urosepsis and demonstrate that thrombocytopenia can be a direct function of bacteremia with UPEC.

## Introduction

Sepsis is a life-threatening organ failure caused by an overactivated immune response to a systemic infection^1^, which, despite improved treatment, is among the most frequent causes of death worldwide.^2^ Thrombocytopenia is generally an independent negative prognostic marker^1,3-5^ and is used as one of the critical parameters to assess the status of critically ill patients in the Sequential Organ Failure Assessment (SOFA) score system. The urinary tract is the primary infection site in around 14% of sepsis cases^6^, often secondary to pyelonephritis. Hence, the Gram-negative, uropathogenic *Escherichia coli* (UPEC)^7^ are frequently implicated and readily cause sepsis-associated thrombocytopenia.^8,9^ Thrombocytes play pivotal roles in infections, contributing to both blood coagulation and immunomodulation (for review, see ^10,11^). The thrombocytopenia that occurs in connection with sepsis is classically associated with a hypercoagulatory state, and *in vitro* clusters of bacteria, such as *Staphylococcus Aureus* and *E. coli*, have been shown to cause local thrombocyte clotting around the bacterial cluster.^12^ While coagulation typically is considered to contain infections, in severe sepsis, this process can lead to multiple organ failure due to diminished blood perfusion and subsequent reduced organ oxygenation.^13,14^ Hence, one would expect that preventing thrombocyte activation and reducing circulating thrombocytes might improve the outcome of sepsis. However, thrombocytes also exhibit an immunomodulatory function by recruiting immune cells for subsequent killing and clearing the infection.^15-20^ Noteworthy, studies support that a reduced number of circulating thrombocytes makes you susceptible to severe infection, and in patients with Primary Immune Thrombocytopenia, a low platelet count correlate to the risk of infection.^21^ This patient group is particularly at higher risk of blood infections^22^ and is at higher risk of severe and fatal infections compared to the general population.^23^ The fall in circulating thrombocytes occurs on the backdrop of the secondary thrombocytosis observed in response to infectious diseases (for review^24^), and several experimental studies suggest that exposure to various bacterial agents or lipopolysaccharide (LPS) results in tissue sequestering of thrombocytes.

In this study, we systematically evaluate the progression of thrombocytopenia in response to UPEC. Our experimental findings present the progression of sepsis in a urosepsis mouse model, helping to understand the consequences of UPEC entry into circulation from an early stage with an initial significant drop in thrombocytes simultaneously with a massive drop in bacterial load. This drop in thrombocytes is followed by a slower reduction associated with activation, thrombus formation, and recruitment of immune cells. Additionally, we detect a significant complex formation between thrombocytes and UPEC that outnumbers thrombocyte-neutrophile complex formation and neutrophile-UPEC complex formation, suggesting a role for thrombocytes as a clearing system in the very early stages of sepsis.

## Methods

### Bacteria

For the experiments, the *E. coli* (UPEC) strain ARD-6 (O6:K13:H1, Statens Serum Institute (Copenhagen, Denmark) as previously characterized^25-27^, was used plain or transfected with enhanced Green Fluorescent Protein (eGFP)-containing plasmid with ampicillin restriction (ARD-6/eGFP-pBAD)^28^. Stocks of UPEC in 15% glycerol Lysogeny Broth (LB)-media (with 100 μg ml^-1^ ampicillin for ARD-6/eGFP-pBAD) were kept at-80°C. New colonies were prepared with 1µl of UPEC stock streaked out on an LB-agar plate with 100 μg ml^-1^ ampicillin for ARD-6/eGFP-pBAD for incubation overnight (37°C) prior to storage at 4°C for up to one month. For overnight liquid cultures, a single colony was transferred to 4 ml LB-medium (with 100 μg ml^-1^ ampicillin for ARD-6/eGFP-pBAD), incubating overnight at 37°C, 250 rpm under aerobic conditions. The overnight cultures were centrifuged (10 minutes, 1300*g*) and washed (5 minutes, 1300*g*), resuspended in sterile saline, and counted by flow cytometry (BD Accuri 6C, BD Biosciences, New Jersey, USA). ARD-6/eGFP-pBAD were prior to washing centrifuged (10 minutes, 1300*g*) and incubated for 3 hours in LB-medium containing 100 μg ml^-1^ ampicillin/0.2% w/v L-arabinose.

### Sepsis model

The experiments were performed in 8-10 weeks old male Balb/CJ mice from Janvier Labs (Saint-Berthevin, France), weighing 24.8±1.5 g, as previously described.^29-32^ The mice were anesthetized by subcutaneous injection of ketamine/xylazine (125 mg kg^-1^/12.5 mg kg^-1^) and placed on a 38°C heating plate. Additional anesthesia (ketamine/xylazine) was administrated according to monitoring of touch-induced movement of whiskers and extremities. The mice received intravenous (iv) bolus injections (150 µl) in one of the tail veins, as previously described,^29,30,32^ containing either sterile saline (0.9% NaCl) or UPEC (ARD-6,165·10^6^/330·10^6^/660·10^6^) dissolved in sterile saline (0.9% NaCl) via a 27G needle. A blood sample was collected from the inferior vena cava into a tube containing sodium citrate (5.5 µmol). The mice were terminated at various time points after the bolus injection, immediately after collecting the blood sample. The time zero represents a termination after the injection and systemic distribution of the UPEC (ca. 10 s corresponding to ∼75 heartbeats). eGFP fluorescent expressing ARD-6/eGFP-pBAD were used for experiments on thrombocyte-UPEC-complex formation. Colony-forming units (CFU) in whole blood were determined by streaking out 50 μl whole blood diluted 2500-fold in sterile 0.9% NaCl solution on LB-agar plates for ImageJ-based assessment after overnight incubation at 37°C. All the experiments were approved by The Animal Experiment Inspectorate, Denmark (2020-15-0201-00422).

### Thrombocyte complex formation

Whole blood was diluted 1:13 in 4-(2-hydroxyethyl)-1-piperazineethanesulfonic acid (HEPES)-buffered salt solution (HBS) and incubated for 30 minutes with the following combinations: *thrombocyte-monocyte complexes*: anti-CD42d-APC (3.83 µg ml^-1^, n°17042180, eBioscience™, Invitrogen)/ anti-Ly6C-FITC (4.78 µg ml^-1^, ab15686, Abcam)/ anti-CD11b-PE (4.78 µg ml^-1^, n°25011281, eBioscience™); *Thrombocyte-neutrophil complexes:* anti-CD42d-APC (3.97 µg ml^-1^, n°17042180, eBioscience™)/anti-Ly6G-FITC (24.8 µg·ml^-1^, n°11966880, eBioscience™). For flow cytometry thrombocytes (APC-positive), monocytes (PE/FITC positive), neutrophils (FITC positive) (Accuri C6 Plus) were counted in samples, which were diluted 1:76 and fixed in Phosphate buffered salt solution (PBS) containing 0.02% formaldehyde. For gating strategies, see Suppl. Fig. 1

### Thrombocyte-UPEC binding

Whole blood collected from mice injected with ARD-6/eGFP-pBAD was diluted 1:2 in 4% paraformaldehyde (PFA) in PBS for 15 minutes, washed twice (5 minutes, 1400*g*), and resuspended in PBS. The sample was diluted 1:5.2 and incubated for 30 minutes with anti-CD42d-APC (3.83 µg ml^-1^, n°17042180, eBioscience™) in the dark. Thereafter, the samples were diluted (1:76, PBS) and thrombocytes (CD42d-positive), UPEC (eGFP-positive), and their aggregations were counted on the flow cytometry (Accuri C6 Plus). For gating strategies, see Fig. 5.

### Whole organ single-cell suspensions

Prior to euthanasia, after blood collection, mice were perfusion fixed through the left ventricle with 4% PFA (5 minutes, 0.2 bar). Subsequently, the brain, kidney, liver, lung, and spleen were isolated and post-fixed in 4% PFA for 10 minutes, followed by washing and storage in PBS (4°C). Each organ is dissected into smaller pieces (∼1 mm^3^) for digestion with 1.5 ml (3 ml for the liver) tissue digestion cocktail for 1 hour (37°C, 50 rpm, shielded from light). Tissue cocktail contains: collagenase/dispase®(1 mg·ml^-1^, Roche n°10269638001), DNase I (0.5 mg·ml^-1^ Fisher Scientific n°10849700) and L-arabinose (100 mg·ml^-1^, Merck n° A91906-1G-A). Digestion was stopped by adding 10 mM EDTA (pH 8), and the preparations were passed through a 70 μm falcon cell strainer (Merck n°CL352350) and rinsed with 25 ml 1% bovine serum albumin (BSA, Sigma, n°A-4503) in PBS. Samples are centrifuged (8 minutes, 350*g*), and the pellets resuspended in PBS. The cells were permeabilized with 0.1% triton x-100 (Sigma-Aldrich X100-100 ml) for 10 minutes and washed with PBS. For staining, cell suspension (10 μl) from each organ was incubated (30 minutes, at RT in the dark) with 3.83 µg ml^-1^ anti-CD42d-APC (n°17042180, eBioscience™, Invitrogen), with 7.66 µg ml^-1^ anti-Mo F4/80 (n°15360960, eBioscience™, Invitrogen) added to the liver samples. Another liver sample was stained with 13.89 µg·ml^-1^ anti-CD105 (n°AF1320, Bio-techne) 30 min prior to staining with anti-CD42d-APC and 34.72 µg·ml^-1^ anti-Goat-IgG-PE (n°AB7004, Abcam) for 30 minutes. The samples were diluted 1:9 and analyzed by flow cytometry (BD Accuri 6C, BD Biosciences). The sample was exited with 488 nm laser for eGFP and PE-Cy7, whereas APC was excited by a 640 nm laser. Emission was detected at 533/30 nm for GFP, 780/60 nm for PE-Cy7, and >670 nm for APC. Data were acquired with the threshold for FSC-H of 10,000, medium speed (35 μl min^-1^). Thrombocytes (APC^+^), Kupffer cells (PE-Cy7^+^), UPEC (eGFP^+^), and complexes of these were counted by flow cytometry (BD Accuri 6C, BD Biosciences). Gating was placed according to unstained samples (See Suppl. Fig. 2).

### Imaging Flow Cytometry

Whole blood samples from mice injected with ARD-6/eGFP-pBAD were centrifuged for 7 minutes at 200*g*. Plasma and buffy coat were collected and further centrifuged for 10 minutes at 1500*g.* The supernatant was removed, and the leukocyte- and thrombocyte-enriched pellet was resuspended in HBS (200 µl). After 30 minutes the preparation was fixed in 4% PFA in PBS for 15 minutes, and washed in PBS (1500*g*, 5 min), and kept overnight at 4°C. The next day, samples were stained for 30 minutes with 4.75 µg ml^-1^ anti-CD42d-APC (n°17042180, eBioscience™, Invitrogen) for detection of thrombocytes together with 7.12 µg ml^-1^ anti-Ly6G-BV421 (n°127627, Nordic Biosite), 3.8 µg ml^-1^ anti-CD11b-PE-Cy7 (n°15370890, Fisher Scientific, Invitrogen) and 7.91 µg ml^-1^ anti-CD115-Alexa Flour 594 (n°135520, Nordic Biosite) for detection of neutrophils and monocytes. Samples were then washed in PBS (1500*g,* 5 min) and resuspended in PBS (50 µl). The samples were analyzed by imaging flow cytometry (ImageStream^®X^Mark II, Luminex, Seattle, WA) equipped with 405 nm, 488nm, 561 nm, and 642nm lasers for relevant excitation. Emission was collected in separate channels with the following filters: Brilliant Violet 421 (435-480 nm), GFP (495-551nm); Alexa 595 (595-625nm); PE-Cy7 (745-780nm); and APC, channel 11 (658-745 nm) and imaged at 60x magnification using INSPIRE^®^ software (version 200.1.620.0, Luminex). The compensation matrix was acquired using INSPIRE^®^ compensation wizard, and matrix calculation and data analysis was performed in IDEAS® software (version 6.3.26.0, Luminex). Spot counting of GFP-fluorescent UPEC in complex with thrombocytes and/or neutrophils was estimated using a GFP mask (Suppl. Fig. 1). Spot counts from neutrophils were evaluated manually to exclude UPEC bound to thrombocytes in complex with neutrophils. The cell area of thrombocytes and neutrophils was determined by masking (Suppl. Fig. 3) and used to determine the number of UPEC per cell area.

### Determination of proinflammatory cytokines

The levels of interleukin-1β (IL-1β), IL-6, keratinocyte chemoattractant (KC, murine equivalent of human IL-8), and tumor necrosis factor-α (TNF-α) in murine plasma were measured by flow cytometry (Accuri C6plus, BD Biosciences) using a CBA flex sets (n°558266, BD Biosciences) according to the manufacturer’s instructions. Plasma was stored for up to 3 months at-20°C prior to measurements.

### Thrombin-antithrombin complex and platelet factor 4

Thrombin-antithrombin complex formation (TAT) was measured by the TAT Complexes Mouse Elisa Kit from Abcam (Cambridge, UK n°ab137994), and Platelet factor-4 (PF-4) was measured by the PF-4 Complexes Mouse Elisa Kit from MERCK (n°RAB0403-1KT) according to manufactures instructions. Plasma was stored for up to 6 months at - 20°C before analysis.

### In vitro bacterial growth

Blood was collected from the inferior vena cava of anesthetized mice in a syringe containing sodium citrate (5 µmol). The samples were centrifuged (400*g*, 10 minutes), and thrombocyte-rich plasma was collected. ARD-6/eGFP-pBAD (8.25 10^6^ ml^-1^) were incubated in ampicillin (100 mg ml^-1^) and 2% arabinose with or without thrombocyte-rich plasma (21.5 μl ml^-1^). The suspensions were incubated at 37°C 250 rpm with sample collection at 0, 30, 60, 90, and 120 minutes. The samples were either diluted 1:100 in PBS for flow cytometer-based growth rate determination or 12.5-fold in 0.9% sterile NaCl solution (50 μl) CFU-determination on LB-agar plates. CFU was counted after overnight incubation (37°C) by ImageJ.

### Bacterial growth and damage in human blood

Human blood was collected from healthy volunteers in EDTA tubes. From blood samples, four different fractions were made: whole blood (WB), platelet-poor plasma (PPP), platelet-rich plasma (PRP), and purified platelets. PRP was made by centrifuging WB (300*g*, 20 minutes without active deceleration), and the supernatant was collected. Platelet-poor plasma was made by centrifuging PRP (2000*g*, 10 minutes, with low deceleration), and the supernatant was collected. The pellet is suspended in PBS to obtain purified platelets. Thrombocytes were stained for 30 minutes: 3.83 µg·ml^-1^ anti-CD42b-APC (n°551061, BD Biosciences). ARD-6/eGFP-pBAD (275.0·10^6^·ml^-1^) and propidium iodide (PI, 150 μM n°P4170, Sigma) were added to 5 different preparations: PBS, WB, PRP, PPP, and purified platelets, and incubated at 37°C, 50 rpm. Samples were collected at various time points over a period of up to two hours. The number of UPEC (eGFP^+^), thrombocytes (CD42b-APC), and PI staining was quantified by flow cytometry (BD Accuri 6C, BD Biosciences). For gating strategies, see Suppl. Fig. 4.

### UPEC and thrombocyte interaction test

Whole blood was fixed 1:1 in 4% PFA in PBS (15 minutes) and washed in PBS (5 minutes, 1500*g*), and the supernatant was replaced with PBS. Various concentrations of anti-CD42b (n°15296947, Invitrogen, Fischer Scientific) were added and incubated for 30 minutes. ARD6/eGFP-pBAD UPEC were added to a final concentration of 165·10^6^ count·ml^-1^ and incubated (10 min, 37°C, 50 rpm, restricted from light). Samples were diluted 1:6.7 in PBS and anti-CD42b-APC (6 μg ml^-1^, n°551061, BD Biosciences) for 30 min (shielded from light). Finally, samples were diluted 1:15.9 in PBS. The number of complexes (GFP/APC^+^) was quantified by flow cytometry (BD Accuri 6C, BD Biosciences).

### Materials

LB broth (Miller, L3522, Sigma), Select agar (n°30391023, Invitrogen). Guanidine buffer: 8M guanidine HCl, 10mM Bis-TRIS, 10 mM dithiothreitol, pH 6. PBS in mM: [Na^+^] 157, [K^+^] 4.5, [Cl^−^] 139.7, [HPO_4_^2-^] 10, [H_2_PO_4_^-^] 1.8, pH 7.4 at 37°C. HBS in mM: [Na^+^] 138, [K^+^] 5.3, [Cl^-^] 132.9, [Mg^2+^] 0.8, [SO_4_^2-^] 0.8, [glucose] 5.6, [HEPES] 14, pH 7.4 at 37°C.

### Statistical analysis

Prior to statistical comparison, the data were tested for outliers and normal/lognormal distribution. A one-way or two-way ANOVA with a Šidák multiple comparisons test was used for normally distributed data according to the experimental design; otherwise, the data were analyzed with Kruskal-Wallis followed by Dunn’s multiple comparisons tests. For statistical differences between time points inside a group, one-way ANOVA with the Šidák multiple comparisons test was used on normally distributed data or Kruskal-Wallis with Dunn’s multiple comparisons test for data not normally distributed. Data were considered statistically significantly different when the adjusted p-values were less than 0.05. Survival studies were analyzed using Kaplan-Meier plots and the Log-rank (Mantel-Cox) test. Data were analyzed using GraphPad Prism 10 software and presented as mean±SEM.

## Results

In our established model for urosepsis^29-31^, we surveyed the development of the systemic response to the presence of circulating UPEC. Adjusting the injection of UPEC (330 million HlyA-producing *E. coli*) to an LD_50_ of approximately 4h (Fig. 1A) allowed us to monitor the development of sepsis and the concomitant change in circulating thrombocytes in anesthetized male mice. To our surprise, the earliest detectable response to UPEC in the bloodstream was a massive reduction of 40.5% in thrombocytes (from 8.212 10^8^±6.069 10^7^ to 4.888 10^8^±9.7869 10^7^, adjusted *p*-value=0.026) in thrombocytes measured as CD42d^+^ cells (Fig. 1B) already after 30 minutes after i.v. introduction of UPEC. This reduced level stabilized over the next few hours, only to show a further drop during the late phases of the infection of up to 63.3% after 4h (adjusted *p*-value<0.0001). Interestingly, the reduction in circulating thrombocytes occurs before any changes in the plasma levels of proinflammatory cytokines can be detected (Suppl. Fig. 5) and was timely associated with a transient increase in thrombocyte activation measured as platelet factor 4 (PF-4, Fig. 1C) and a corresponding transient increase in thrombin anti-thrombin (TAT)-complex formation 30 minutes after injection of the bacteria (Fig. 1D). Of note, both PF-4 and TAT complexes stabilized at levels similar to controls not exposed to bacteria, long before systemic signs of intravascular coagulation. Hence, the coagulation-induced use of thrombocytes in micro-thrombosis is an unlikely explanation for the acute reduction in thrombocyte count.

**Figure 1.**
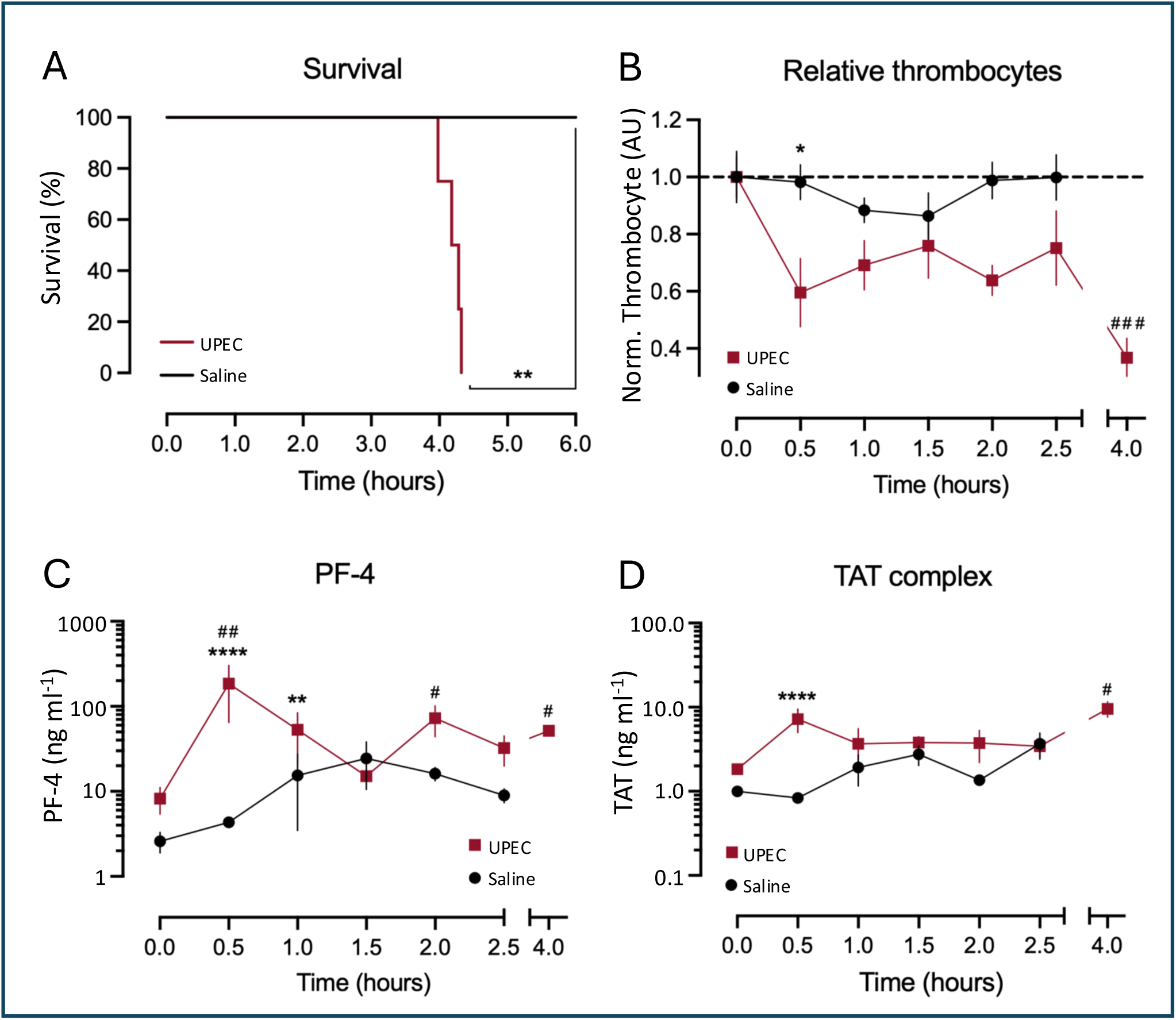
Bacteraemia-induced reduction in circulating thrombocytes and relation to coagulation and thrombocyte activation. **(A)** Kaplan-Meier plot showing survival of male mice over 6 h after exposure to 330·10^6^ UPEC (sepsis) or control (saline), n=4 for each survival-study group. **(B)** shows the number of circulating CD42d^+^ thrombocytes in mice terminated at various time points after *iv*-injection of 330·10^6^ UPEC. **(C)** shows ELISA-determined plasma concentrations of PF-4 and **(D)** TAT complexes. Data are given as mean±SEM, n=4-7 for each group \**p*<0.05, \*\**p*<0.01, \*\*\**p*<0.001, \*\*\*\**p*<0.0001.

Another option for the apparent reduction in circulating thrombocytes is that thrombocytes form complexes with recruited leukocytes. Generally, thrombocytes are known to form complexes with neutrophils^33^ and monocytes ^34,35^ during inflammation and sepsis. Figure 2 shows that mice exposed to UPEC i.v. showed a biphasic pattern with an initial reduction in circulating Ly6G^+^ neutrophils (Fig. 2A) after 30 minutes, followed by an increase that peaked after two hours. Interestingly, the neutrophil thrombocyte complexes follow the same pattern, with an early reduction within the first 30 minutes and a subsequent increase (Fig. 2B). Since the level of neutrophils that do not bind thrombocytes remains stable (Fig. 2C), the early reduction in neutrophils results from a reduction in neutrophil-thrombocyte complexes. Gating neutrophils for maturity shows that the early sepsis-induced reduction in neutrophils represents a removal of mature neutrophils (Fig. 2D and E). In contrast, the later increase represents the recruitment of immature neutrophils into the circulation, potentially secondary to the rise in the plasma concentration of PF-4. Since thrombocytes are known to guide leukocytes to sites of extravasation^36^, the reduction of mature neutrophils (from 3.404 10^6^±1.253 10^6^ at time zero to 0.364±0.089 10^6^ after 30 minutes) could potentially contribute to the early thrombocytopenia. Assuming 1:1 binding of thrombocyte to neutrophils, this would only account for 0.9% of the thrombocyte reduction, and 10:1 only 9%. Considering Ly6C^+^/CD11b^+^ monocytes, there is a tendency for an early reduction in circulating monocytes, but this is not statistically significant. In the later phases, monocytes are also recruited into the blood (Fig. 2F), and the monocyte-thrombocyte complexes follow the same pattern (Fig. 2G), whereas the unbound monocyte level remains stable (Fig. 2H). Hence, the acute 40.5% drop in circulating thrombocytes after induced bacteremia cannot be explained by mere adhesion to circulating leukocytes. However, the number of circulating mature neutrophils is reduced in parallel with the reduction in circulating thrombocytes.

**Figure 2.**
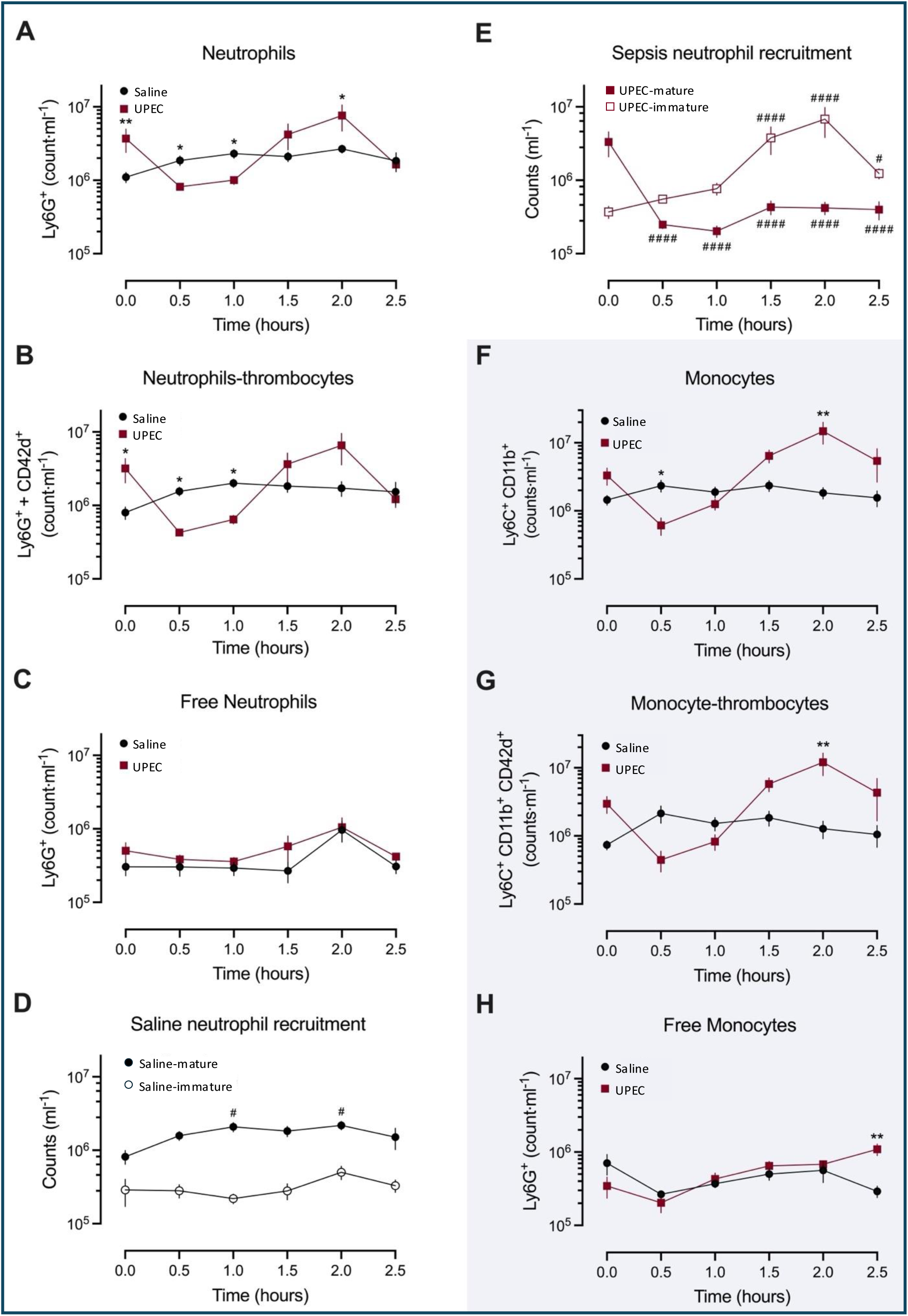
Recruiting leucocytes and complexes formation with thrombocytes during sepsis. Whole blood was collected from male mice at various time points after i.v.-injection of 330·10^6^ ARD-6. **(A)** shows the number of circulating neutrophils (Ly6G^+^), **(B)** the number of neutrophil-thrombocyte aggregates Ly6G^+^/CD42d^+^, and **(C)** neutrophils that do not bind thrombocytes. **(D)** illustrates the number of mature and immature neutrophils Ly6G^+^ in saline controls. **(E)** shows that change in immature/mature neutrophils after i.v.-injection of 330·10^6^ ARD-6. **(F)** represent the corresponding change in total Ly6C^+^, CD11b^+^ monocytes, **(G)** Ly6C^+^, CD11b^+^/CD42d^+^ monocyte-thrombocyte aggregates, and **(H)** monocytes that do not bind thrombocytes by flow cytometry. Data are given as mean±SEM, n=4-7 for each group \**p*<0.05, \*\**p*<0.01, \*\*\**p*<0.001, \*\*\*\**p*<0.0001. The full statistics for C and D are included in Suppl. Tabel 1

The early thrombocytopenia and reduction in mature neutrophils are timely associated with clearance of injected UPEC. Figure 3A shows that the bacterial content in a blood sample collected from mice ∼10 s/75 heartbeats after injection of UPECs falls drastically within the first 30 minutes and remained constantly low for the remaining observation period. This acute drop indicates that the UPEC are either removed from the circulation, sticking to the endothelium, or neutralized in the circulation. Based on this finding, we speculated that the thrombocytes may play an essential function in the acute clearance of UPEC in the circulation. To address this aspect, we introduced an eGFP-expressing plasmid in our ARD-6 strain (ARD-6/eGFP-pBAD), and Figure 3B shows imaging flow cytometry using gating masks for GFP^+^, CD42d^+^, and Ly6G^+^ cells. Strikingly, Figure 3B demonstrates that ARD-6/eGFP-pBAD immediately binds to the thrombocytes and neutrophils after injection of UPEC into the blood (Fig. 3B). We find evidence for UPEC binding to single thrombocyte, single neutrophil, thrombocytes in aggregates, and neutrophil-thrombocyte complexes. Figure 4 shows the binding of UPEC to thrombocytes and neutrophils, which occurs immediately after injection of the UPEC. Thrombocytes have the largest capacity for binding UPEC at any timepoint, going from 77.9% of all UPEC bound to thrombocytes, singlet or in aggregates, at time zero to 98.2% after 10 minutes. In contrast, neutrophils account for 1.7% and 1.8%, respectively (Fig. 4A). Figure 4B illustrates most thrombocytes bind only one bacterium, whereas neutrophils generally bind more bacteria per cell. However, Figure 4C demonstrates that thrombocytes appear to be less efficient in binding UPEC, demonstrating a significantly lower number of UPEC bound per unit surface area for thrombocytes compared to neutrophils. Thus, it is primarily the abundance of the thrombocytes that potentially makes them a relevant clearance system for invading UPEC.

**Figure 3.**
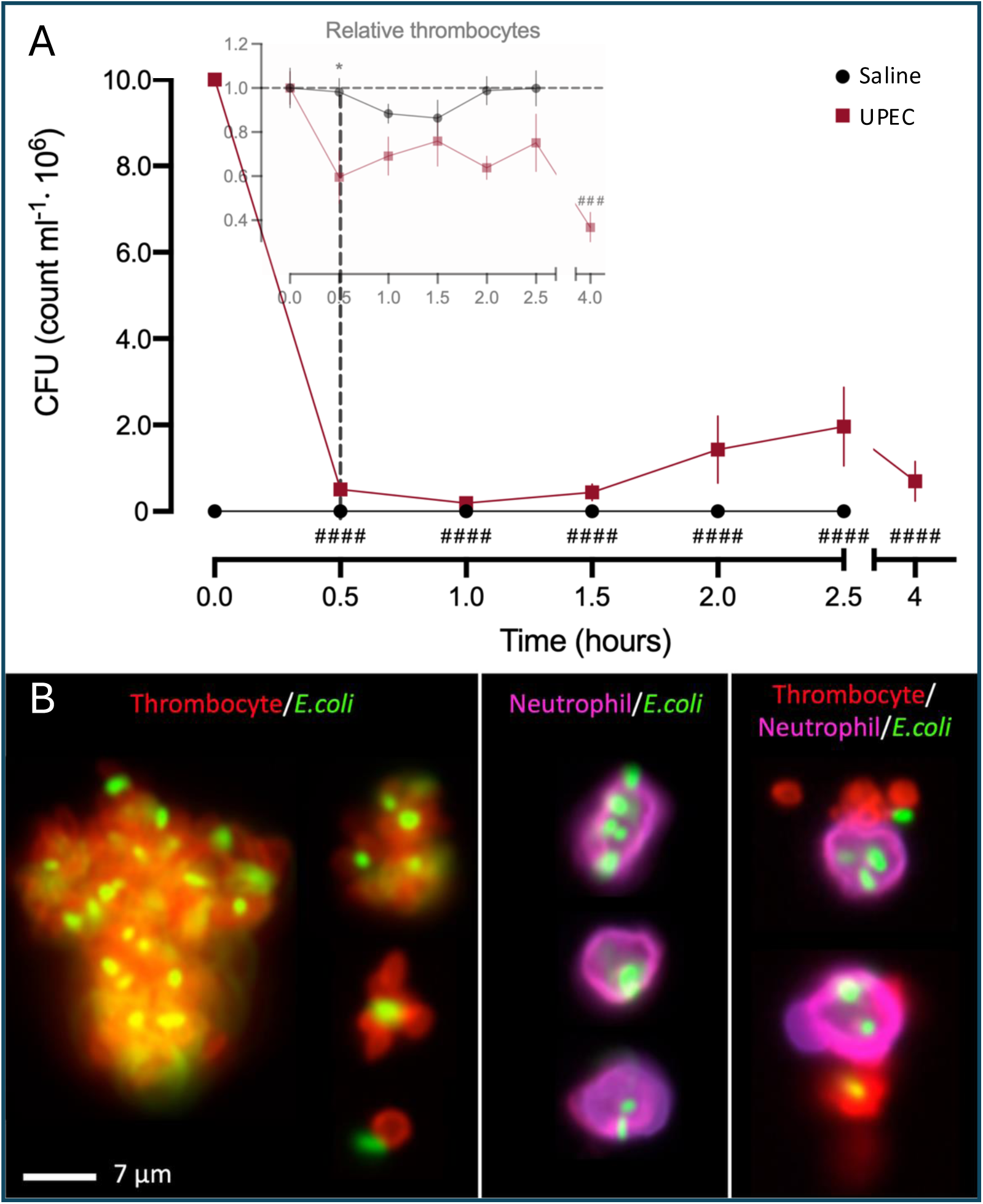
Injected UPEC bind to thrombocytes and are immediately removed from the circulation. **(A)** Bacterial load in whole blood was determined as colony-forming units at various time points after i.v.-injection of 330·10^6^ ARD-6 in male mice, n=4-7 for each group. Insert is a comparison with Fig. 1B. Data are given as mean±SEM, * represents *p*<0.05 for comparison saline vs. UPEC, whereas ^###^ represents *p*<0.001 and ^####^ *p*<0.0001 compared to time zero. **(B)** Representative images from imaging flow cytometry of *E. coli* complex formation: CD42d-APC (thrombocytes, red), ARD-6/eGFP-pBAD (*E. coli,* green), CD11b-PE-cy7/Ly6G-BV421 (neutrophils, pink/purple) stained blood collected at time zero (∼20 s) after injection of *UPEC*.

**Figure 4.**
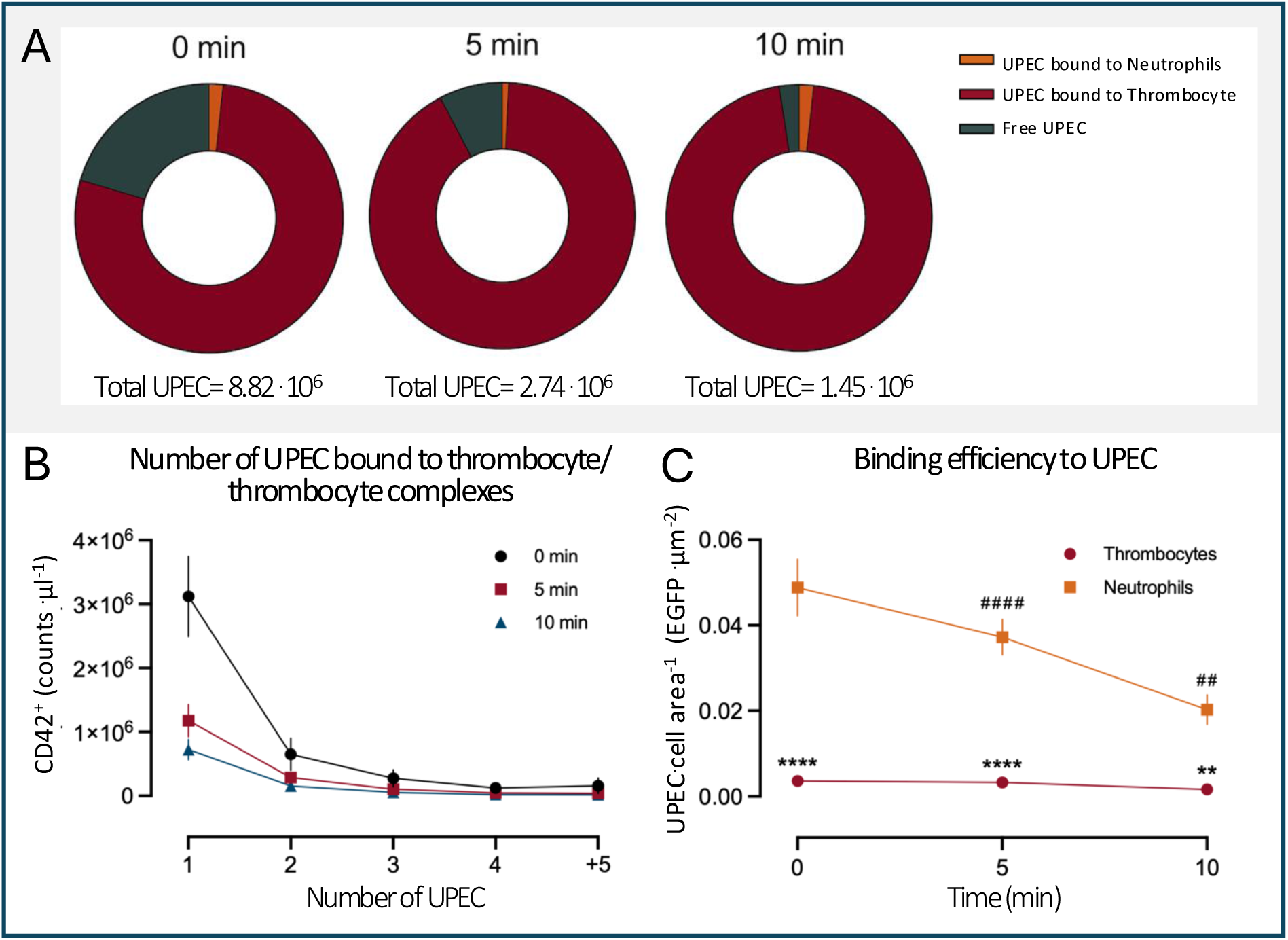
Thrombocytes constitute a low affinity, high-capacity clearance system for UPEC. Blood was collected at various time points (0, 5, or 10 minutes) after i.v.-injection of 330·10^6^ ARD-6/eGFP-pBAD in male mice. Imaging flow cytometry was used for data acquisition. **(A)** shows pie charts of the total amount of UPEC (eGFP^+^) and the percentage bound to thrombocytes (CD42d^+^) and neutrophils (Ly6G^+^) at each time point. **(B)** shows the results of a GFP mask used to estimate the number of eGFP^+^ UPEC bound to each thrombocyte complex. **(C)** illustrates UPEC per cell area (neutrophil or thrombocyte) based on area masks combined with an eGFP-spot-count mask. *n*=3-4 for each group. Data are given as mean±SEM. ** marks *p*<0.01, \*\*\*\**p*<0.0001 for comparison between groups and ^##^*p*<0.01, ^####^ *p*<0.0001 for comparison to time zero for neutrophils.

Because UPEC is bound to thrombocytes instantaneously, we zoomed in on the observation period from 0-10 minutes after UPEC administration. Figure 5A-C shows the gating for ARD-6/eGFP-pBAD, CD42^+^, and complexes. Figure 5D shows that the UPEC is cleared rapidly (91%) within the first 10 minutes. This clearance did not affect the number of thrombocytes, which does not bind bacteria (Fig. 5E). However, there is a clear reduction in the thrombocyte-UPEC complexes in the same period. Given that the distribution between thrombocytes carrying more than one UPEC and the fraction of thrombocyte-bound bacteria are similar to what is seen in the imaging flow cytometry (ca. 53% after 0 minutes and 100% after 10 minutes), with the fraction increasing with time, we conclude that the majority of the bacteria are cleared in complex with thrombocytes.

If thrombocytes are to be an efficient scavenging system for UPEC, their capacity must be overridden to affect the effective bacterial load. Therefore, we challenged the thrombocyte binding capacity with various UPEC concentrations to learn if it is possible to overload the system (Fig. 6A). Interestingly, at a dose of 165 10^6^ UPEC, we did not observe any reduction in circulating thrombocytes over the first hour of observation. However, doubling the initial dose to 660 10^6^ UPEC resulted in no further reduction in thrombocyte count (Fig 6B). Correspondingly, we show that the effective number of circulating bacteria was similarly low at a dose of 165 10^6^ and 330·10^6^ UPEC and increased dramatically at a UPEC dose of 660 10^6^. A cautious interpretation of this observation is that overloading the thrombocyte’s capacity for binding UPEC results in delayed bacterial clearance. Our data support the notion that the binding of UPEC to thrombocytes may be involved in the clearance of UPEC from circulation. With most of the thrombocytes binding UPEC 1:1, one would expect the thrombocytes and UPEC to disappear from circulation in quantitatively equal amounts. To test this, we directly counted the number of UPEC in circulation using our ARD-6/eGFP-pBAD as a more accurate measure for bacterial clearance than CFU. Figure 5D shows an early reduction in circulating UPEC (8.2·10^7^, when assuming a blood volume for mice of 1 ml). Unfortunately, the reduction in total circulating thrombocytes did not become statistically significant ten minutes after UPEC injection. However, if we allow ourselves to calculate the difference in mean value at time zero (5.83 10^8^ cells/ml) and after 10 minutes (4.61 10^8^ cells/ml), the discrepancy is 12.2 10^7^ cells ml^-1^ (Fig. 5E). This number is in the vicinity of the reduction seen in circulating UPEC, particularly considering the tendency for thrombocytes to aggregate following the activation seen upon UPEC exposure. Simultaneously, we also observe a reduction in thrombocyte-UPEC complexes (Fig. 5F). However, it is not possible to make a proper estimation of disappeared complexes because they are constantly formed and removed during the process.

**Figure 5.**
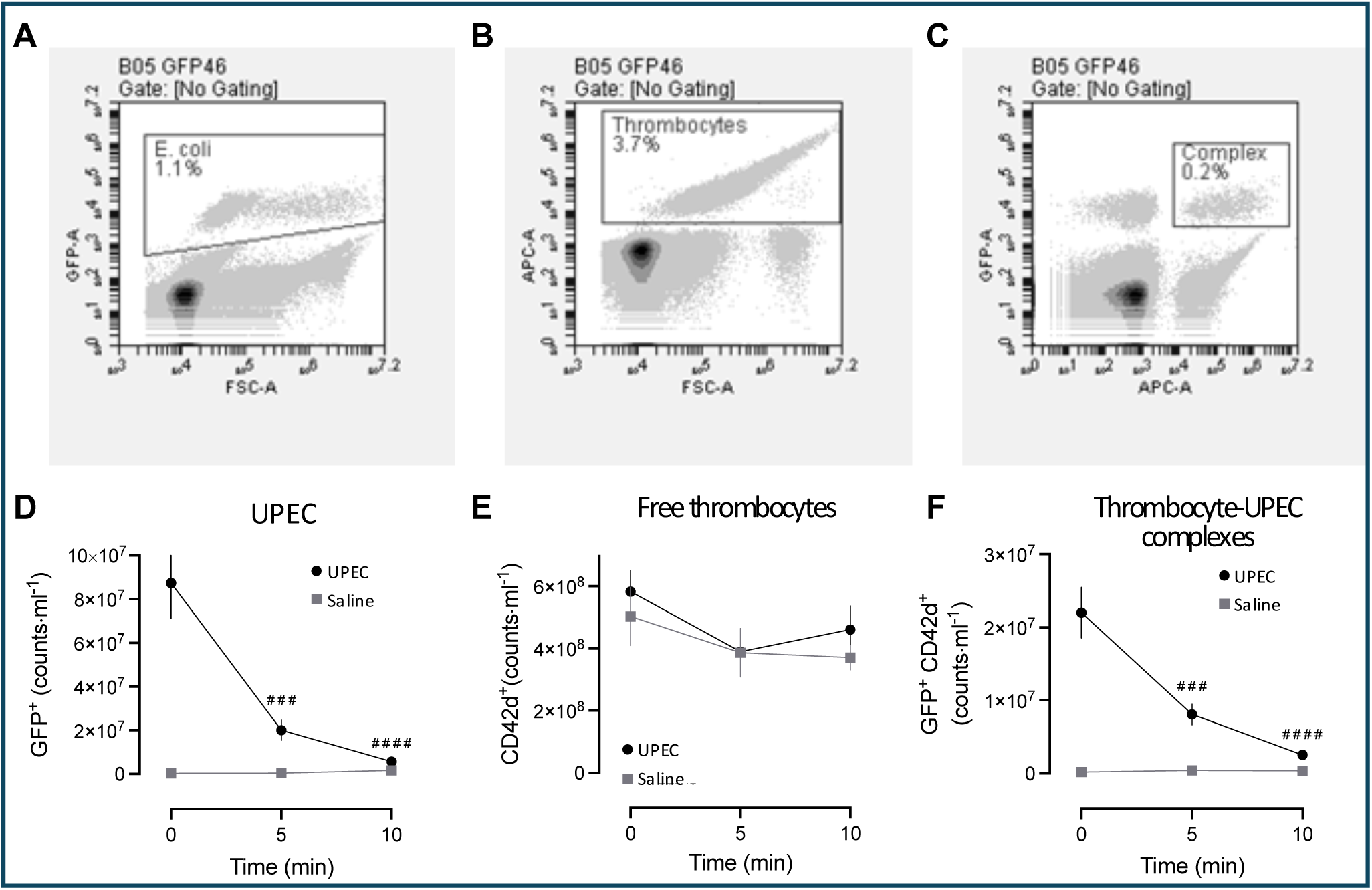
Instantly formed *thrombocyte-UPEC* complexes are cleared from the bloodstream. Whole blood was collected after different incubation periods (0, 5 or 10 minutes) of bacteremia induced with i.v.-injection of 330·10^6^ ARD-6/eGFP-pBAD in male mice. Plots with a gating strategy from flow cytometry are shown in **(A)** for eGFP^+^ UPEC as eGFP-A, and **(B)** for CD42d^+^ thrombocytes (APC-A) and **(C)** for complexes positive for both UPEC and thrombocyte markers. For all time points, **(D)** eGFP^+^ UPEC, **(E)** CD42d^+^ thrombocytes and **(F)** eGFP^+^- and CD42d^+^-complexes were quantified. *n*=6 for each group. Data are given as mean±SEM, ^###^*p*<0.001, ^####^*p*<0.0001 compared to time zero in the UPEC group.

**Figure 6.**
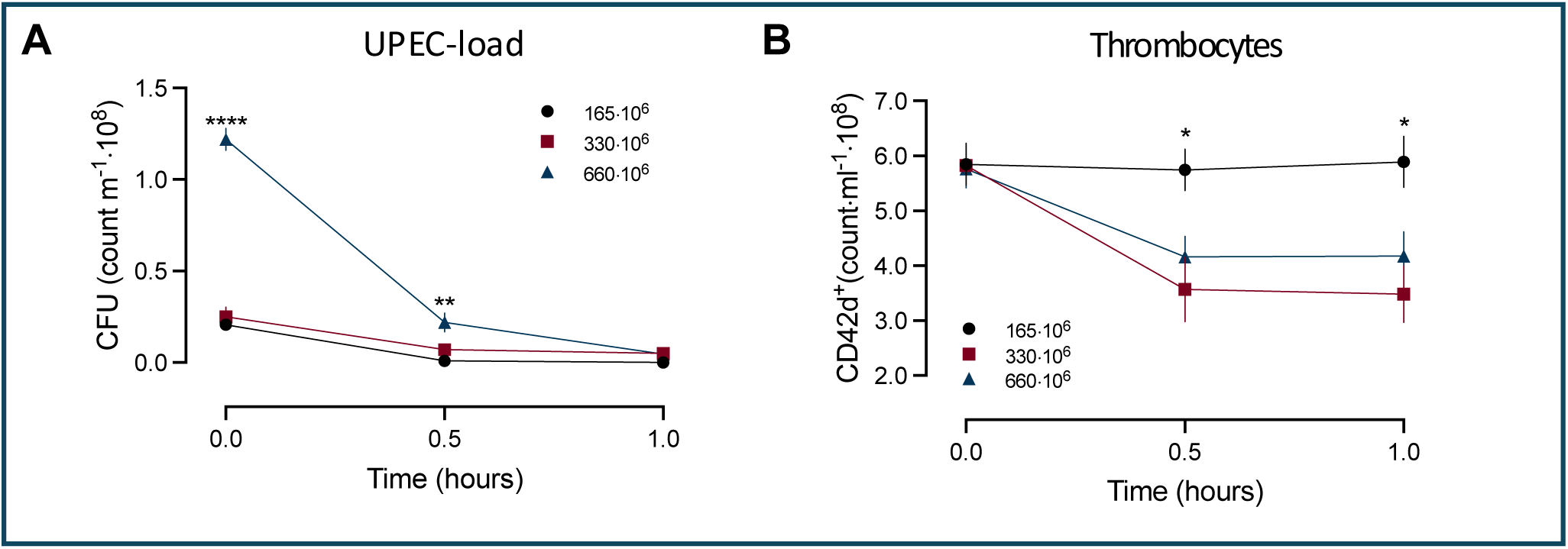
Thrombocytes’ capacity for UPEC clearance. Blood was collected at various time points (0, 30 min, 60 min) after i.v.-injection of 165·10^6^, 330·10^6^ or 660·10^6^ ARD-6 in male mice. **(A)** Bacterial load determined as colony-forming units. **(B)** show the parallel changes in circulating thrombocytes (CD42d^+^, flow cytometry). *n*=4-7 for each group. Data are given as mean±SEM. * indicates a comparison of UPEC (165·106) and either of the other two groups (UPEC 330·10^6^ and UPEC 660·10^6^). \**p*<0.05, \*\*\*\**p*<0.0001

Moreover, we determined the acute tissue distribution of UPEC as a measure of the acute clearance of circulating bacteria. We used mice exposed to ARD-6/eGFP-pBAD or saline for 0 or 30 minutes, followed by perfusion fixation. The data showed that CD42^+^ thrombocytes in the saline controls are found mainly in the spleen and liver. Upon exposure to UPEC, the proportion of thrombocytes immediately increases in the liver and continues for the first 30 minutes after the introduction of UPEC to the bloodstream, whereas the thrombocyte count in the other organs remains constant (Fig. 7A). We confirm our previous finding that the tissue accumulation of thrombocytes is paralleled by a fall in circulating thrombocytes (Fig. 7B). Simultaneously, we observe a massive accumulation of the ARD-6/eGFP-pBAD in the liver, with very little exposure to the other organs (Fig. 7C), which follows the clearance of UPEC from the circulation (Fig. 7D). Please note that practically equal amounts of UPEC present at time zero in the circulation is found in the liver at time 30 minutes (Fig. 7C and D). The picture is quite similar for the thrombocyte/UPEC complexes, with a main accumulation in the liver, although some of the complexes seemingly accumulate in the lung (Fig. 7E). Again, the tissue accumulation mirrors the clearance of complexes from the circulation (Fig. 7F).

**Figure 7.**
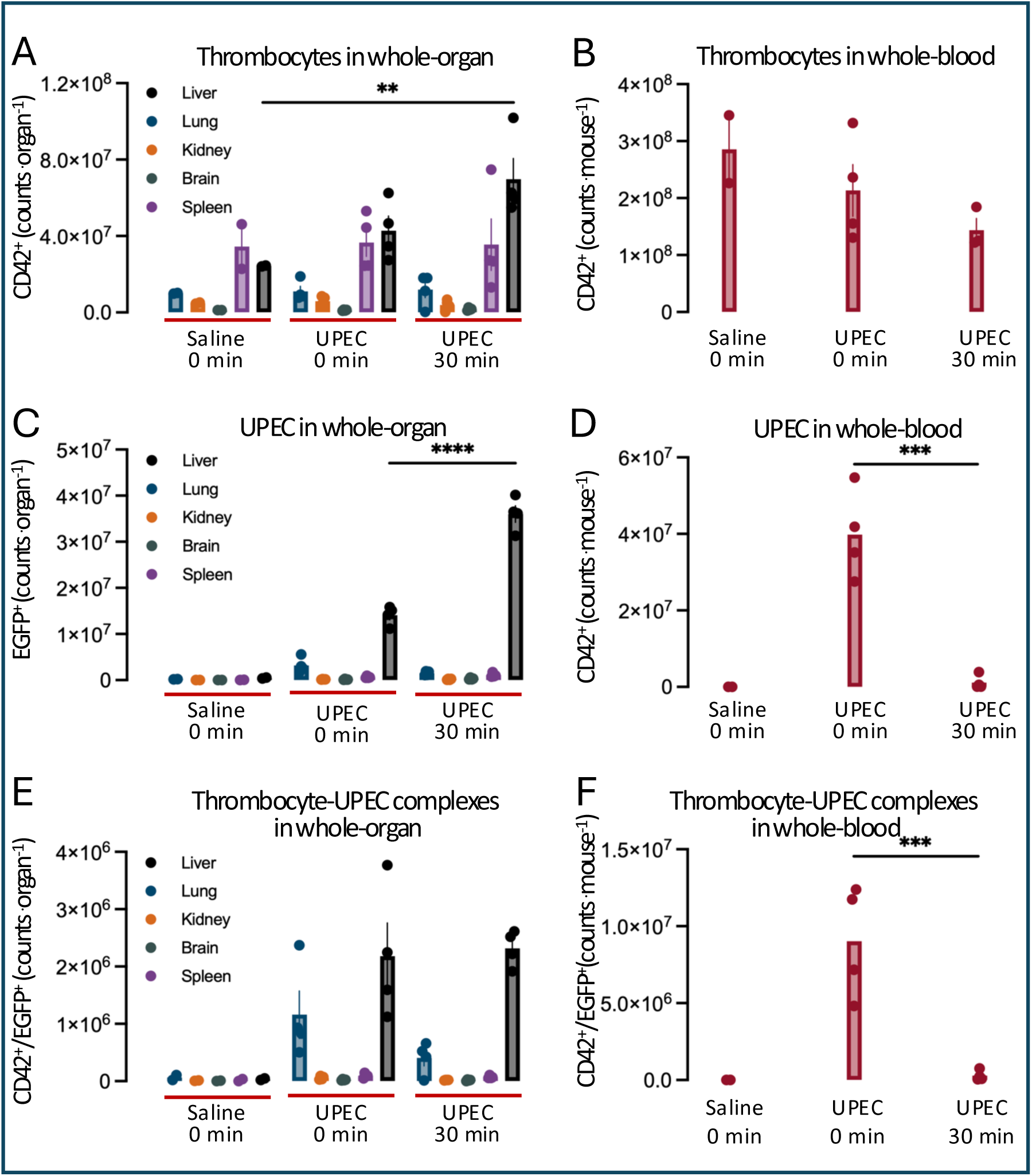
Time-dependent tissue accumulation of thrombocytes, UPEC and their complexes. Single-cell suspensions of PFA-perfused whole organs or whole blood 0- or 30-minutes post i.v.*-*injections of either saline or 330·10^6^ ARD-6/eGFP-pBAD in male mice. **(A)** shows the total number of thrombocytes (CD42d^+^) in the liver, lung, kidney, brain, and spleen and **(B)** the corresponding change in total CD42d^+^ in the circulation. **(C)** illustrates the total UPEC (eGFP^+^) number of the liver, lung, kidney, brain, and spleen, and **(D)** the parallel circulating eGFP^+^ UPEC. **(E)** shows all thrombocyte/UPEC complexes in the liver, lung, kidney, brain, and spleen, and **(F)** the corresponding total thrombocyte/UPEC complexes in the circulation. Data are given as mean±SEM, \**p*<0.05, \*\**p*<0.01.

Our data support that the vast majority of the injected UPEC is sequestered in the tissue. However, several studies suggest that thrombocytes might be directly bactericidal through the release of antimicrobial peptides^37^. Opposed, UPEC could potentially be responsible for reducing thrombocyte numbers by directly cytotoxic effects on the thrombocytes^31^. Surprisingly, we could not detect any effect of adding thrombocytes on the growth of ARD-6/eGFP-pBAD in LB medium over two hours of incubation measured either by flow cytometry or as CFU (Fig. 8A and B). Since thrombocytes are activated and even lysed by pore-forming bacterial toxins like α-hemolysin (HlyA) produced by UPEC^31^, one could speculate that the sudden reduction in thrombocyte count could result in platelet damage by HlyA. However, the presence of UPEC does not seem to have any influence on the number of thrombocytes over a 2h incubation, and hence, the fall in circulating thrombocytes cannot be explained by cytolysis (Fig. 8C). This finding is supported by intravascular hemolysis, which in our model is shown to be a functional assessment of HlyA-production by UPEC^38^, occurring long after the initial drop in circulating thrombocytes. Regardless of preparation, the thrombocyte number is stable over time, and only a minor fraction (up to 0.15%) shows damage over 2h measured as propidium iodide positive (PI^+^) cells (Fig. 8E). We do, however, see a substantial reduction in UPEC over 2h incubation in preparations where plasma is present, with around 30% PI^+^ after 2h in platelet-rich and platelet-poor plasma (Fig. 8D and E). These data underscore that, most likely, complement activation with major attack complex formation and antimicrobial peptides in the plasma will affect the UPEC content of the blood over time. Notably, the fraction of PI^+^ UPEC is 0.027 (CI 95% = -0.002; 0.056) after 30 minutes and can only account for a minor fraction of the early bacterial clearance. This is underscored by the fact that the majority of the UPEC are recovered by the liver within the first 30 minutes. Moreover, the early reduction of thrombocytes in response to bacteremia is not caused by bacteria-induced thrombocyte damage.

**Figure 8.**
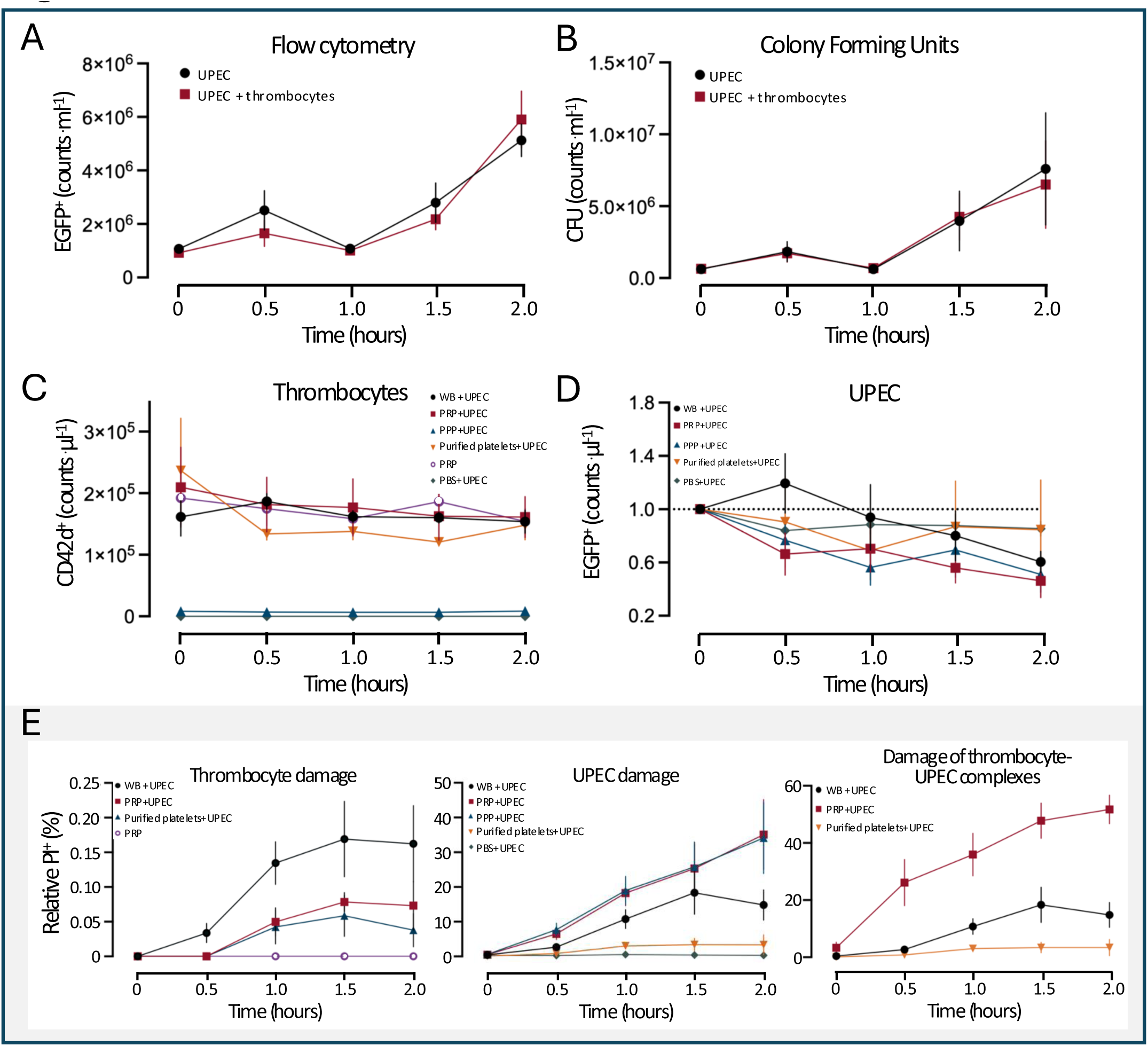
Thrombocytes do not inhibit bacterial growth in vitro. UPEC growth in LB-media with or without thrombocytes. **(A)** shows the effect of thrombocytes on the number of UPEC (eGFP^+^) over time (0-2 hours) counted by flow cytometry and **(B)** the number of CFU. *n*=2-3 for each group. **(C)** depicts the change in the total number of thrombocytes during *in vitro* incubation of whole blood (WB), platelet-rich plasma (PRP), platelet pure plasma (PPP), and purified platelets in the absence or presence of UPEC, and **(D)** shows the parallel changes in total UPEC. **(E)** illustrates the corresponding changes in a number of damaged thrombocytes, UPEC, and thrombocyte/UPEC complexes over time, where cell damage is determined by a propidium iodide assay (PI). *n*=3-5 for each group. Data are given as mean±SEM. Statistics can be found in Suppl. Table 2.

Capture and clearance of infectious agents in the liver, in general, occurs through a duad of the sinusoidal endothelial cells for smaller particles (<0.3 µm) and for larger particles, the phagocytotic Kupffer cells (for review, see^39^). Surprisingly, only a small fraction of thrombocytes and UPEC at any given time is co-localizing with CD11^+^ used as a marker for Kupffer cells, which we expected to effectuate the UPEC clearance (Suppl. Fig. 6). In the liver, UPEC are found unbound or in complex with thrombocytes, with only a minor fraction adhering to Kupffer cells either at time 0 or after 30 minutes. However, the fraction of UPEC that colocalized with endothelial cells either alone or in complex with thrombocytes was detected at time zero, and the fraction increased over the first 30 minutes (Suppl. Fig. 6A). It must be emphasized that the majority of the UPEC are found unattached to either of the cell types at any of the timepoints. For the thrombocytes, a similar fraction colocalizes with Kupffer cells both at time zero and after 30 minutes. Similarly, we see a fraction of thrombocytes that adhere to endothelial cells either alone or in complex with UPEC (Suppl. Fig 6B). Correspondingly, the fraction of thrombocyte-UPEC complexes is reduced over time. The thrombocytes were also found predominantly to be unbound to either of the cell types in the liver. Overall, this suggests that UPEC are not phagocytosed to a large extent by the Kupffer cells. There is a transient attachment to the endothelial cells, and the thrombocyte-UPEC complex formation is not sustained upon hepatic clearance.

## Discussion

Thrombocytopenia is an integrated part of the body’s response to bacteremia and a hallmark of the septic state as a validated biomarker for negative outcome.^1,3-5^ Sepsis-induced thrombocytopenia has long been primarily ascribed to a hypercoagulation state secondary to intense pressure on pro-inflammatory pathways.^40^ However, recently, experimental studies have emphasized that thrombocyte activation, complex formation with various leukocytes, and tissue sequestration contribute to the thrombocytopenia observed during sepsis (for review, see ^41^). This could potentially explain why thrombocytopenia in itself is a risk factor for severe infections.^21,23^ In this study, we systematically investigated the development of thrombocytopenia after introducing UPEC into the blood as a model of a rapidly developing urosepsis. Our model allows us to separate intravascular coagulation from other factors that might reduce the number of circulating thrombocytes as a function of time. We detected an acute 40.5% reduction in single circulating platelets already 30 minutes after UPEC had been administered i.v. The dramatic, acute fall in platelets could not be explained by either intravascular coagulation, intravascular lysis, or damage of the thrombocytes. Despite thrombocytes being known to form complexes with neutrophils^33^ and monocytes^34,35^ and, through complex formation, guide leukocytes to inflammatory sites for extravasation^36^, early removal of mature neutrophils can only explain a fraction of the reduction in thrombocytes during the first 30 minutes of observation.

Clearly, the acute reduction in circulating thrombocytes was accompanied by a marked reduction in circulating UPEC. We show that the injected UPEC immediately binds to the thrombocytes and neutrophils in the circulation and that thrombocytes, through their sheer number, were able to bind around 80% of the UPEC immediately after they were added to the blood, whereas neutrophils bind less than 2%. We demonstrate that the injected UPEC are efficiently cleared from the circulation within the first 30 minutes and that the early reduction of the thrombocytes primarily occurs in those that have formed complexes with UPEC. Moreover, we observe cluster formation between thrombocytes that bind UPEC, but the majority of UPEC are in 1:1 complex with thrombocytes. We show that the early reduction of circulating thrombocytes is a function of the bacterial load and that when the binding capacity for UPEC on thrombocytes is surpassed, the number of free UPEC in the blood remains high over time.

Our data suggest that thrombocyte-UPEC complexes are cleared in the tissue, and our data on tissue sequestering support that this primarily happens in the liver. A substantial part of the thrombocytes that are reduced in response to UPEC in the blood goes to the liver, most likely for degradation. This coincides with the fact that the majority of the UPEC are sequestered in the liver and that only a minor fraction is seen by other tissues. Collectively, our data support that thrombocytes are essential for effective bacterial clearance and that a high bacterial load can overwhelm the thrombocyte binding capacity, resulting in a higher number of free, circulating, uncleared bacteria.

Interestingly, there are reports supporting a close association between the introduction of bacteria in the bloodstream and a drastic fall in circulating thrombocytes. A Canadian study from 1976 shows that cases of accidentally contaminated infusion resulted in an acute drop in circulating thrombocytes without signs of intravascular coagulation and similar thrombocyte reductions during other acute infections that preceded the increase in circulating neutrophils.^42^ This is supported by experimental studies using LPS to inflict a sepsis-like response, where acute injection of LPS causes a reduction of circulating thrombocytes^43^ as a result of platelet aggregation and adhesion to endothelial cells^44^. Interestingly, mice expressing an interleukin-4 receptor/ GPIbα fusion protein that lacks GPIb-IX receptor function showed better survival compared to wildtype mice after exposure to i.v. LPS and blocking GPIb-IX receptor function improved the survival after LPS exposure^44^. These data are supported by a recent study showing that deletion of the GPIb-IX cytoplasmic tail better the survival of *Staphylococcus aureus* and *E. coli*-sepsis with reduced sequestering of thrombocytes in the liver in response to *S. aureus^45^.* This is surprising in the light of our data, showing the importance of platelets for UPEC clearance, since one would expect that reducing tissue sequestering of thrombocytes will increase the UPEC load in the blood. Interestingly, the c-terminal deficient GPIb-IX mice showed reduced clearance of S. *aureus^45^,* and we showed a similar finding on UPEC clearance by preventing thrombocyte activation and thrombocytopenia with a P2Y_1_ receptor antagonist^31^. This illustrates that preventing platelet activation and sequestration might be hazardous in the treatment of bacterial sepsis. In support of this notion, it has been shown that experimental depletion of thrombocytes reduced the survival and bacterial load of *Bacillus cereus*^46^, *Klebsiella pneumoniae*^47^, and UPEC^32^ in circulation. Hence, our data strongly suggest that platelets are sequestered in the tissue as part of the bacterial clearance of thrombocyte/bacterial complexes, at least in the case of UPEC.

Several studies support that invading bacteria directly activate thrombocytes, and the ability to activate and aggregate thrombocytes is a specific trait of certain bacteria, such as the Gram-positive *Staphylococcus aureus^48,49^* and *Bacillus cereus^46^*. This potentially suggests that bacteria-induced thrombocytopenia is linked to specific strains causing severe sepsis. Here, we show that UPEC instantaneously binds to thrombocytes immediately upon introduction into the blood and that the binding is timely associated with a transient increase in thrombocyte activation, i.e., PF-4 secretion. We show that the reduction in circulating thrombocytes is a direct function of the UPEC load, and the reduction in thrombocytes is a consequence of the removal of thrombocyte-UPEC complexes from circulation. The notion that platelets are essential gatekeepers of systemic infections is supported by a cardinal paper from 2017 on thrombocyte migration, showing that thrombocytes bind, trap, and inactivate fibrin-coated *E. coli* in the circulation.^18^ Our data show that thrombocytes bind UPEC directly in the circulation and underscore that if the capacity for entry point screening of bacteria is overridden, direct binding to thrombocytes in the bloodstream can support elimination.

We fully acknowledge that bacteria are rarely infused in substantial amounts directly into the bloodstream, as is the case in our model. However, our model allows us to isolate one of the fundamental mechanisms in response to bacterial infection and quantify its capacity. We show that practically all infused UPEC are recovered in the liver and that the liver’s capacity for clearing UPEC is fast and effective, leaving only a fraction of the UPEC to be incapacitated through antimicrobial peptides or complement in plasma. Interestingly, the UPEC bound to thrombocytes are not substantially affected by the potential release of antimicrobial peptides in response to bacterial-induced thrombocyte activation since only plasma-containing preparations and not isolated thrombocytes affect UPEC growth *in vitro*.

Our findings reciprocate the notion of an effective clearance system for circulating bacteria in the liver, which is generally recognized for its importance in the clearance of bloodborne microbes. The hepatic clearance of infectious agents mainly occurs in the sinusoids, constituting a two-cell type system, where the smaller particles (<0.3µm) are cleared by the sinusoidal endothelial cells and the larger ones by the phagocytotic Kupffer cells (for review, see^39^). Kupffer cells have previously been shown to clear circulating bacteria in a thrombocyte-dependent manner.^46^ However, in this elegant study, thrombocytes were shown to have ‘touch and go’ interactions with Kupffer cells, but after binding of *B. anthracis* to the Kupffer cells, the thrombocytes altered behavior and adhered in a GPIIb-dependent manner.^46^ We find UPEC to bind to thrombocytes directly in the circulation immediately after injection, and these thrombocyte-UPEC complexes are readily cleared. We cannot completely exclude that part of the thrombocytopenia observed is secondary to the primary binding of UPEC to Kupffer cells with subsequent thrombocyte adhesion. However, we only observe a small subfraction of UPEC to interact with Kupffer cells, either alone or in complex with thrombocytes. One potential explanation could be that either the UPEC capsule or complex formation with thrombocytes prevents binding to the Kupffer cells. In this context, an important study on thrombocytes’ role in bacterial clearance suggests that thrombocyte-bacterial interaction prevents immediate bacterial clearance^50^. The study provides evidence for *Listeria monocytogenes* being cleared by a dual-track system, the fast track being Kupffer cell-dependent clearance of free bacteria in the blood, and the slow-path requiring C3-dependent opsonization for binding to thrombocytes. The latter will spare the bacteria from destruction in the liver and allow presentation for splenic dendritic cells.^50^ The study provides data that implies that the same system applies to Gram-negative *E. coli*. This finding is in sharp contrast to our current data. Notably, we do not find C3 to be important for UPEC’s interaction with thrombocytes because we can detect substantial thrombocyte interaction between purified thrombocytes and UPEC, and antiCD42, even in high concentrations, did not prohibit thrombocyte-UPEC interaction (Suppl. Fig. 7). The discrepancy is likely to lie in the choice of *E. coli* strain since the capsular K polysaccharides markedly reduce *E. coli* strains’ ability to bind C3.^51^ Hence, ATCC 25922 (O6:H1)^51^ may potentially be optimized with C3, whereas this is not the case for intact UPEC, as used in this study (O6:H1:K13). Therefore, the described dual-track clearance does not apply for urosepsis, where direct thrombocyte interaction is important for instantaneous clearance of UPEC in a C3-independent fashion.

We do, however, find the UPEC/thrombocyte-UPEC complexes to interact with the sinusoidal endothelial cells. This suggests that this cell type is likely to be involved in the clearance of UPEC from the circulation. There are previous reports of commensal *E. coli* from the gastrointestinal tract being able to bind to sinusoidal endothelial cells and, via flagella interaction with TLR5, augment fibrosis in Non-alcoholic Fatty Liver Disease^52^. Since not all UPEC are found to adhere to endothelial cells, one can speculate that it is only a transient adhesion before extravasation. Thrombocytes have previously been implicated in the extravasation of cancer cells during metastasis^53^ and also in the liver sinusoidal endothelial cells^54^, and one could speculate that thrombocytes are essential for the opening of the endothelial barrier that would allow UPEC to pass between the cells into the interstitium (for review, see^55^). Whether this mechanism has any relevance for the clearance of UPEC has to be determined.

In conclusion, our data support that thrombocytes constitute an effective, high-capacity, low-affinity system for acute clearance of UPEC that invade the circulation. We demonstrate that GFP-expressing UPEC is bound instantaneously to thrombocytes in the circulation and that the clearance of thrombocyte-UPEC complexes is associated with a marked drop in circulating thrombocytes. These findings might also explain the potential benefit of secondary thrombocytosis frequently associated with infection, with *E. coli* as one of the frequent agents^56^. Importantly, a high circulating bacterial load can cause thrombocytopenia, and hence, a low thrombocyte count can potentially reflect a high bacterial load. This could be an important contributing factor for thrombocytopenia being a negative prognostic marker in sepsis.

## Acknowledgments

Large thanks must be given to Lab manager Helle Jakobsen for skilled technical support. Imaging flow cytometry was performed at the FACS core Facility, Aarhus University, Denmark. Funding was received from Novo Nordisk Foundation NFF 0064953 and NNF21SA0069371.

## Authorship Contributions

Nanna Johnsen (NJ), Mette H. Christensen (MC), and Helle Praetorius (HP) have founded the concept for study. NJ has conducted the majority of the experiments with contribution from MC and Laura V. Sparsø (LVS).

Emil H. Lambertsen has contributed with experiment for the supplementary data, and Thomas Corydon (TC) has produced GFP-expressing UPEC and contributed to the finalization of the manuscript.

