## Supplementary Material for "Platelet-dependent clearance of uropathogenic *Escherichia coli* directly drives sepsis-induced thrombocytopenia in a mouse model"

### Supplementary figure 1

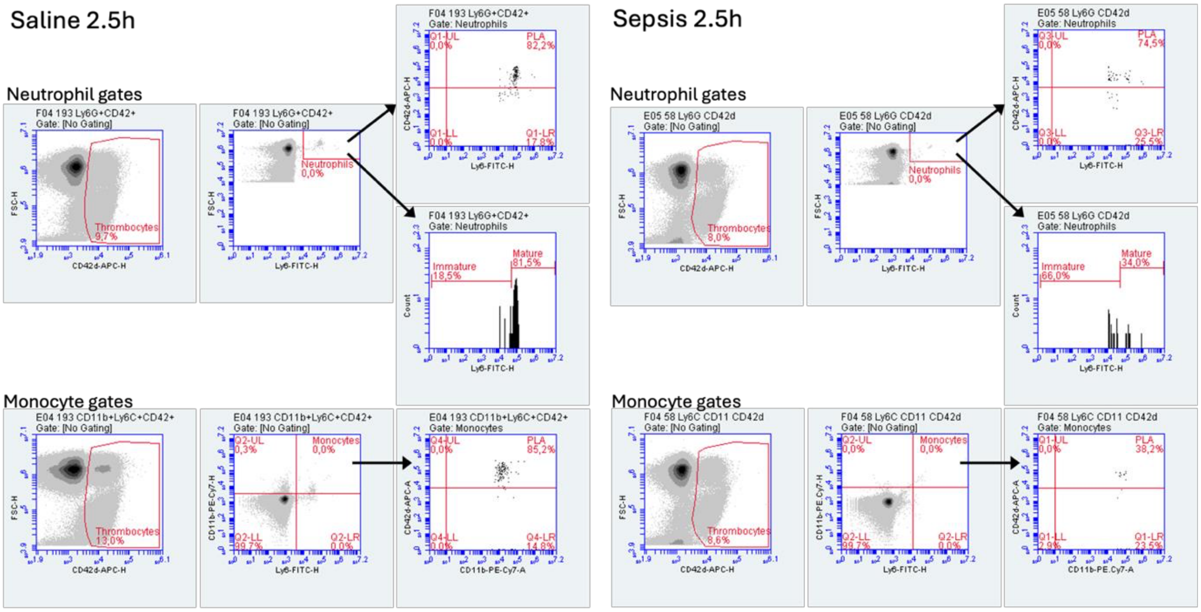

***Suppl. Fig. 1 – Gating strategies for Thrombocyte-Leukocyte aggregations in murine blood samples*** *Flow cytometry gating strategy on scatterplots showing representative data (2.5h saline and 2.5h sepsis). anti-CD42d-APC was used to identify thrombocytes. Anti-Ly6C-FITC identify neutrophils. Based on the neutrophil gate, neutrophil-thrombocyte complexes and neutrophil maturation were identified. Antibodies recognizing CD11b and Ly6C conjugated with FITC and PE-Cy7 respectively, were used to identify monocytes. Based on the monocyte gate, monocyte-thrombocyte complexes were identified.*

### Supplementary figure 2

A

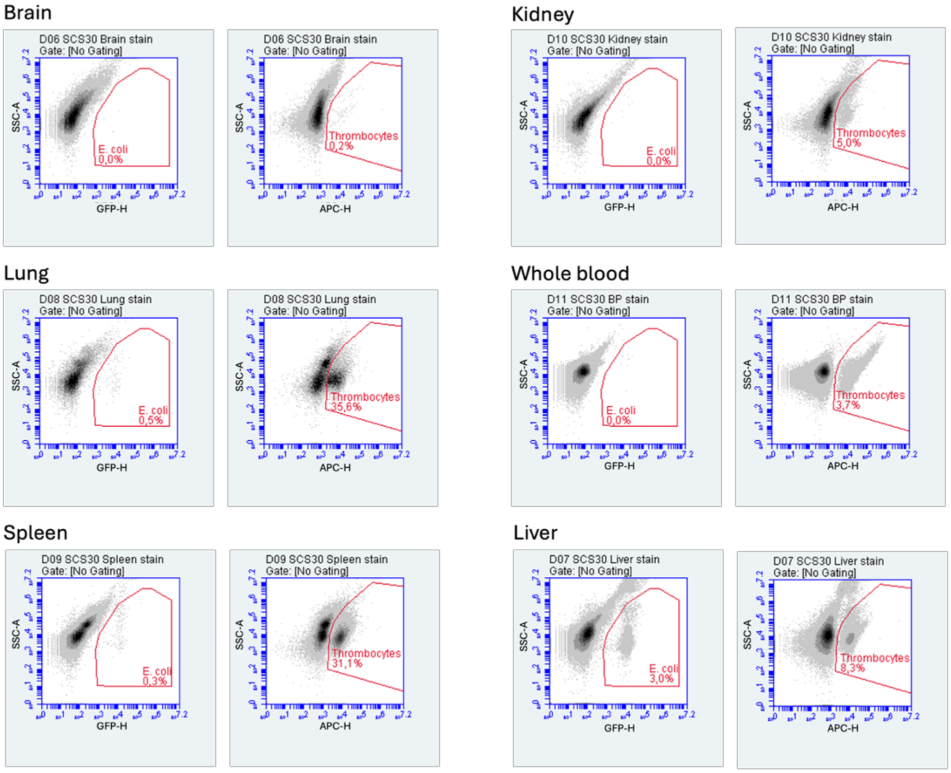

| Gate \ Population | $GFP^+$,$CD42^+$ |
|---|---|
| *E. coli* | Intersection |
| Thrombocytes | Intersection |

B

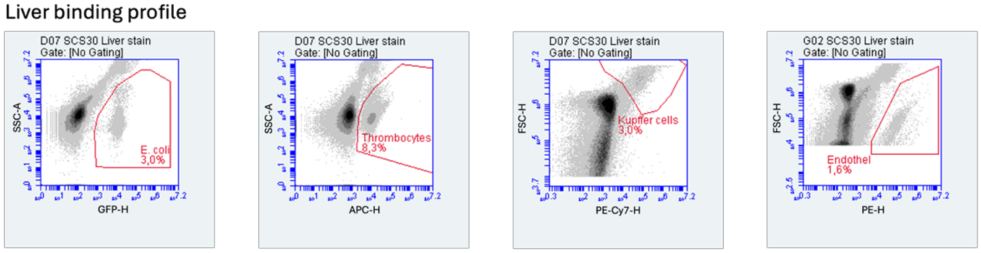

| Gate \ Population | $GFP^+$,$CD42^+$,$F4/80^-$ | $GFP^+$,$CD42^-$,$F4/80^+$ | $GFP^-$,$CD42^+$,$F4/80^+$ | $GFP^+$,$CD42^+$,$F4/80^+$ | $GFP^+$,$CD42^-$,$CD105^+$ | $GFP^-$,$CD4^+$, $CD105^+$ |
|---|---|---|---|---|---|---|
| *E. coli* | Intersection | Intersection | Remove | Intersection | Intersection | Remove |
| Thrombocytes | Intersection | Remove | Intersection | Intersection | Remove | Intersection |
| Kupffer cells | Remove | Intersection | Intersection | Intersection | NA | NA |
| Endothel | NA | NA | NA | NA | Intersection | Intersection |

*Suppl. Fig. 2 – Gating strategies for organ single cell suspension Flow cytometry gating strategy on scatterplots showing representative data (30 min septic mouse). (**A**) anti-CD42d-APC was used to identify thrombocytes. GFP to identify E. coli. Table shows how complexes was determined by intersection of E. coli and thrombocytes gates. (**B**) anti-CD42d-APC was used to identify thrombocytes. GFP to identify E. coli. anti-F4/80-PE-Cy7 identifies kupffer cells and anti-CD105-PE identifies endothelial cells. Different combination was used to quantify different populations (se table).*

### Supplementary figure 3

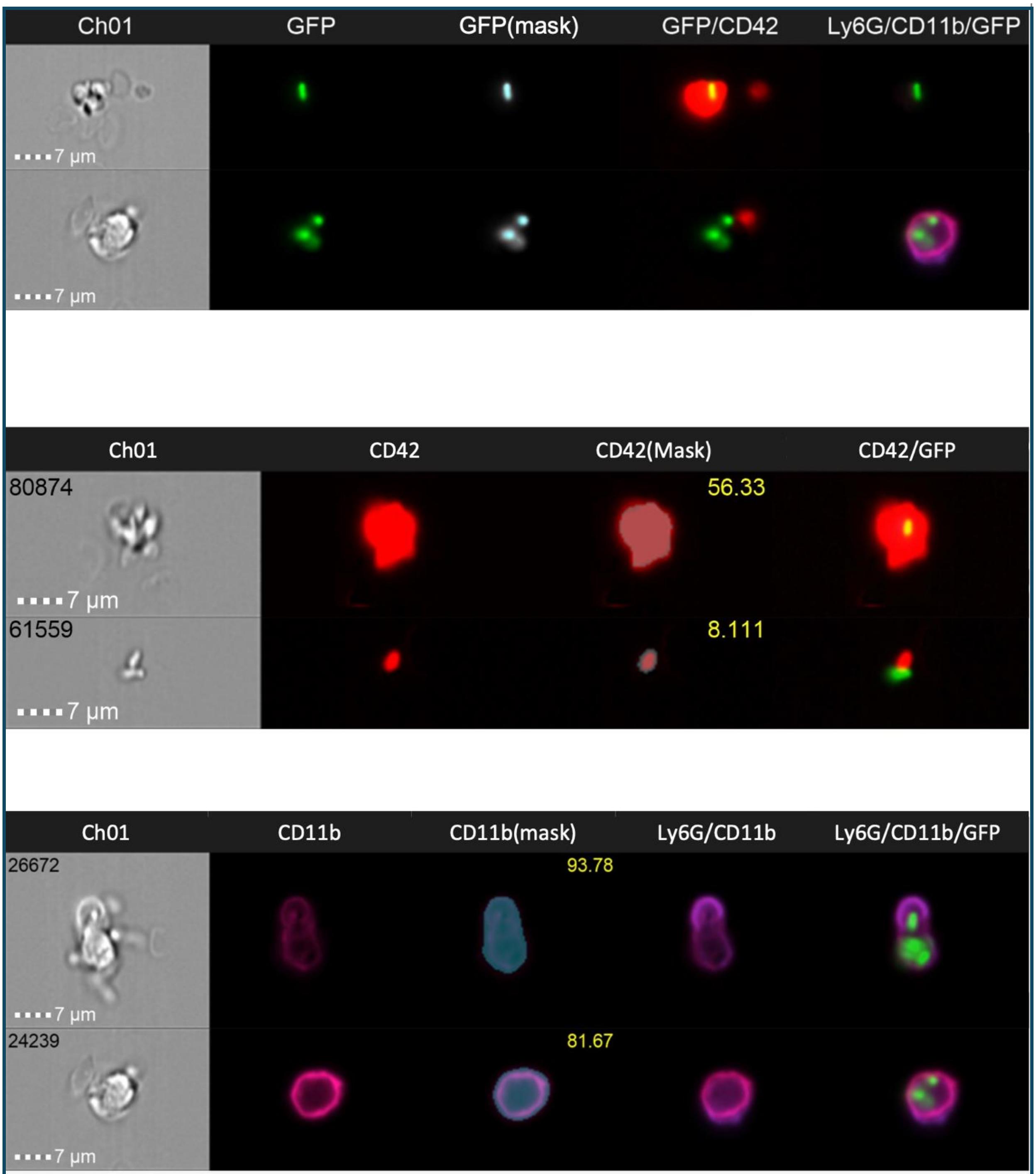

*Suppl. Fig. 3 - Mask strategies for imaging flow cytometry. The 3 panels show masking strategies for E. coli (GFP) spot counting, CD42d (thrombocyte) area and CD11b (neutrophil) area.*

### Supplementary figure 4

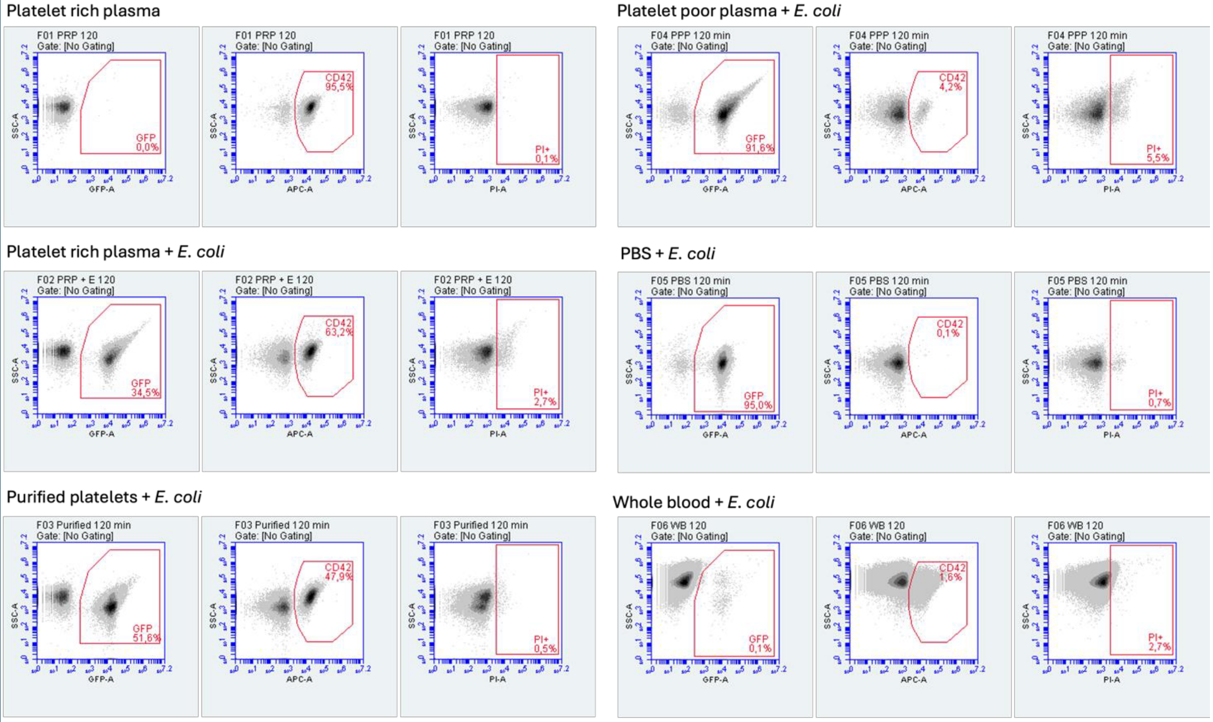

***Suppl. Fig. 4 – Gating strategies for bacterial growth and damage*** *Flow cytometry gating strategy on scatterplots showing representative data (2.0h). anti-CD42d-APC was used to identify thrombocytes. GFP to identify E. coli and propidium iodide (PI) to identify damaged thrombocytes and/or E. coli. Intersection of GFP and CD42 gate was used to identify complexes and intersection with PI-gate was used to determine damage of each population.*

### Supplementary figure 5

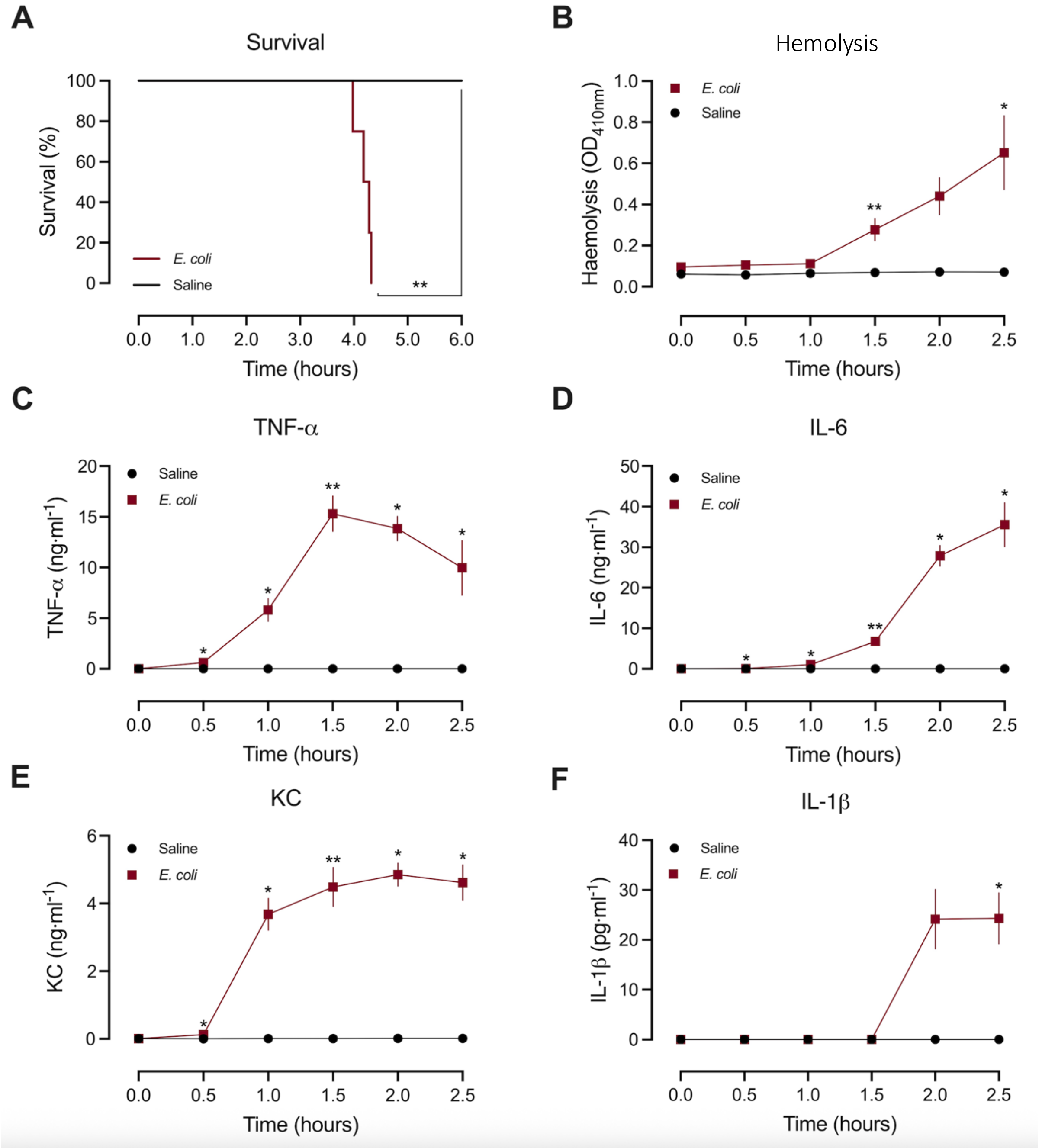

*Suppl. Fig. 5 – Characteristics of the septic model.* *Bacteraemia was induced with iv-injection of UPEC (330·$10^6$ ARD-6) in male mice. (**A**) Kaplan-Meier plot shows survival over 6h after exposure to HlyA-producing UPEC or control (saline). n=4 for each survival-study group. Blood was collected after various incubation periods (0, 0.5, 1.0, 1.5, 2.0, and 2.5 h) of bacteraemia to study the progression of sepsis. (**B**) Progression of haemolysis measured on plasma ($OD_{410}$nm). Flow cytometry CBA flex set was used to determine the progression in plasma levels of proinflammatory cytokine (**C**) TNF-α, (**D**) IL-6, (**E**) KC and (**F**) IL-1β. n=4-7 for each group. Data are given as mean ±SEM, *p<0.05, **p<0.01, ***p<0.001, ****p<0.0001.*

### Supplementary figure 6

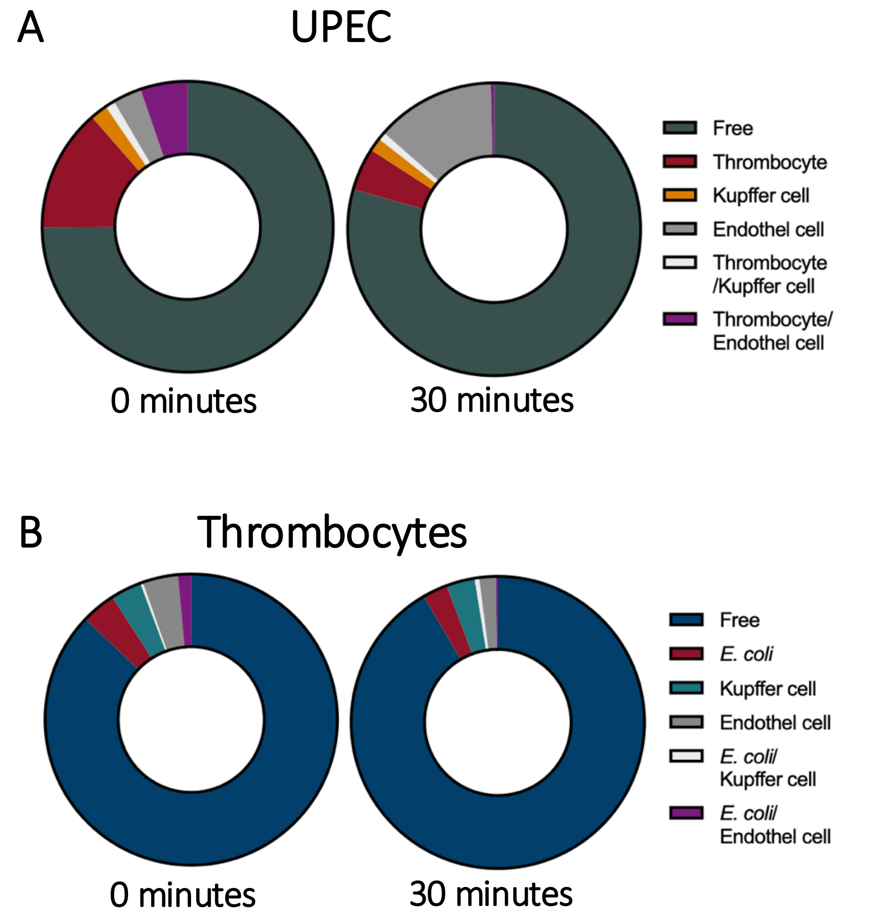

***Suppl. Fig. 6 – Hepatic clearance of UPEC and thrombocytes – association to the sinusoidal clearance system.*** *(**A**) Pie chart of all UPEC in the liver, alone (eGFP$^+$), UPEC thrombocyte complexes alone (eGFP$^+$/CD42$^+$), UPEC adhesion to Kupffer cells alone (eGFP$^+$/CD11$^+$), triple positive (eGFP$^+$/CD42$^+$/CD11$^+$), and UPEC adhesion to sinusoidal endothelial cells (eGFP+/CD31+) and triple positive (EGFP$^+$/CD42$^+$/CD31$^+$). (**B**) Pie chart of total thrombocytes in the liver, alone (eGFP$^+$), thrombocytes in complexes with UPEC alone (CD42$^+$/eGFP$^+$), thrombocyte adhesion to Kupffer cells alone (CD42$^+$/CD11$^+$), triple positive (CD42$^+$/eGFP$^+$/CD11$^+$), and thrombocyte adhesion to sinusoidal endothelial cells (CD42$^+$/CD31+) and triple positive (CD42$^+$/ EGFP$^+$/CD31$^+$). n=2-4 for each group.*

### Supplementary figure 7

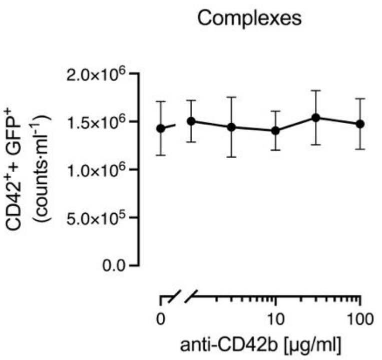

***Suppl. Fig. 7 – Complex formation is not prevented by blocking the CD42b receptor.*** *Fixated whole blood was incubated with various concentration of anti-CD42b. Number of GFP- and CD42d-positive complexes were counted. n=4. Data are given as mean ±SEM.*

### Supplementary table 1

| | Saline, mature | Saline, immature | Sepsis, mature | Sepsis, immature |
|---|---|---|---|---|
| 0.0 vs. 0.5 | 0.4458 | 0.9096 | **<0.0001** | 0.2550 |
| 0.0 vs. 1.0 | 0.1456 | 0.9967 | **<0.0001** | 0.1234 |
| 0.0 vs. 1.5 | 0.4023 | **0.0097** | **0.0002** | **0.0037** |
| 0.0 vs. 2.0 | **0.0426** | 0.9966 | **<0.0001** | **<0.0001** |
| 0.0 vs. 2.5 | 0.9417 | **0.0217** | 0.2571 | 0.8652 |

*Suppl. Table 1 – Statistics for Figure 2C and D*

### Supplementary table 2

| | | Adjusted P-values | | | | | |
|---|---|---|---|---|---|---|---|
| | | WB + *E. coli* | PRP + *E. coli* | Purified platelets + *E. coli* | PPP + *E. coli* | PBS + *E. coli* | PRP |
| Thrombocyte | 0.0 vs. 0.5 | 0.9077 | 0.568 | 0.671 | 0.5995 | 0.7434 | 0.9352 |
| | 0.0 vs. 1.0 | >0.9999 | 0.4077 | 0.7054 | 0.4829 | >0.9999 | 0.5845 |
| | 0.0 vs. 1.5 | >0.9999 | 0.3943 | 0.5842 | 0.4099 | 0.7434 | 0.9994 |
| | 0.0 vs. 2.0 | 0.9988 | 0.4905 | 0.7229 | 0.9985 | 0.7434 | 0.473 |
| Thrombocyte Death | 0.0 vs. 0.5 | 0.6158 | >0.9999 | >0.9999 | | | >0.9999 |
| | 0.0 vs. 1.0 | **<0.0001** | 0.3593 | 0.5059 | | | >0.9999 |
| | 0.0 vs. 1.5 | **<0.0001** | 0.0626 | 0.2872 | | | >0.9999 |
| | 0.0 vs. 2.0 | **<0.0001** | 0.0919 | 0.5999 | | | >0.9999 |
| *E. coli* | 0.0 vs. 0.5 | 0.8025 | 0.2447 | 0.995 | 0.2865 | 0.2007 | |
| | 0.0 vs. 1.0 | 0.9965 | 0.3919 | 0.5398 | 0.1153 | 0.4475 | |
| | 0.0 vs. 1.5 | 0.6985 | **0.0472** | 0.9826 | 0.4794 | 0.5242 | |
| | 0.0 vs. 2.0 | **0.0384** | **0.0344** | 0.9774 | 0.0907 | 0.1854 | |
| *E. coli* Death | 0.0 vs. 0.5 | 0.3088 | 0.1121 | 0.1507 | 0.0979 | >0.9999 | |
| | 0.0 vs. 1.0 | 0.0589 | **0.0491** | 0.169 | 0.0606 | 0.8975 | |
| | 0.0 vs. 1.5 | 0.12 | 0.1058 | 0.3959 | 0.1056 | >0.9999 | |
| | 0.0 vs. 2.0 | 0.1127 | 0.1052 | 0.6948 | 0.1209 | 0.9949 | |
| Complex | 0.0 vs. 0.5 | 0.5094 | 0.9768 | 0.6971 | | | |
| | 0.0 vs. 1.0 | 0.3908 | 0.4214 | 0.6275 | | | |
| | 0.0 vs. 1.5 | 0.3898 | 0.696 | 0.6043 | | | |
| | 0.0 vs. 2.0 | 0.231 | 0.7805 | 0.5643 | | | |
| Complex Death | 0.0 vs. 0.5 | 0.3088 | 0.0968 | 0.1507 | | | |
| | 0.0 vs. 1.0 | 0.0589 | **0.027** | 0.169 | | | |
| | 0.0 vs. 1.5 | 0.12 | **0.0221** | 0.3959 | | | |
| | 0.0 vs. 2.0 | 0.1127 | **0.0114** | 0.6948 | | | |

*Suppl. Table 2 – Statistics for Figure 7*
